# FruitFire: a luciferase based on a fruit fly metabolic enzyme

**DOI:** 10.1101/2023.06.30.547126

**Authors:** Spencer T. Adams, Jacqueto Zephyr, Markus F. Bohn, Celia A. Schiffer, Stephen C. Miller

## Abstract

Firefly luciferase is homologous to fatty acyl-CoA synthetases from insects that are not bioluminescent. Here, we determined the crystal structure of the fruit fly fatty acyl-CoA synthetase CG6178 to 2.5 Å. Based on this structure, we mutated a steric protrusion in the active site to create the artificial luciferase FruitFire, which prefers the synthetic luciferin CycLuc2 to D-luciferin by >1000-fold. FruitFire enabled in vivo bioluminescence imaging in the brains of mice using the pro-luciferin CycLuc2-amide. The conversion of a fruit fly enzyme into a luciferase capable of in vivo imaging underscores the potential for bioluminescence with a range of adenylating enzymes from nonluminescent organisms, and the possibilities for application-focused design of enzyme-substrate pairs.

## Introduction

Fruit flies, like most insects, do not glow in the dark. But why not? The fruit fly *Drosophila melanogaster* expresses a long-chain fatty acyl-CoA synthetase (ACSL), CG6178, which is 41% identical to *Photinus pyralis* firefly luciferase (Fluc).^1^ Insect ACSLs are promiscuous enzymes, recognizing a variety of fatty acid substrates.^2^ Substrate promiscuity leading to the adenylation of the Fluc substrate D-luciferin could potentially allow subsequent oxidation and light emission (Figure 1), and has been hypothesized to underlie the evolution of beetle luciferases from ACSLs.^3–5^ However, unlike fireflies and other bioluminescent beetles, nonluminescent insects such as fruit flies do not biosynthesize D-luciferin, which is required for light emission in beetles (Figure 1).^6^ Moreover, despite its homology to Fluc (Figure 2), CG6178 has been shown to be incapable of adenylating D-luciferin, and does not emit light when treated with D-luciferin and ATP.^1,2^ Nonetheless, further characterization of CG6178 has revealed that the enzyme is a latent luciferase capable of catalyzing light emission, not from D-luciferin, but from synthetic luciferin analogues such as CycLuc2.^7,8^ Although this latent luciferase activity is relatively weak, it suggests that fatty acyl-CoA synthetases can inform us about how beetle luciferases evolved from their evolutionary antecedents. Further, employing these concepts to engineer enzymes like CG6178 may yield novel and selective luciferases for challenging *in vivo* imaging applications.^9–12^ We therefore sought to understand the molecular basis for this substrate discrimination and to improve the luciferase activity of this ACSL enzyme.

**Figure 1.**
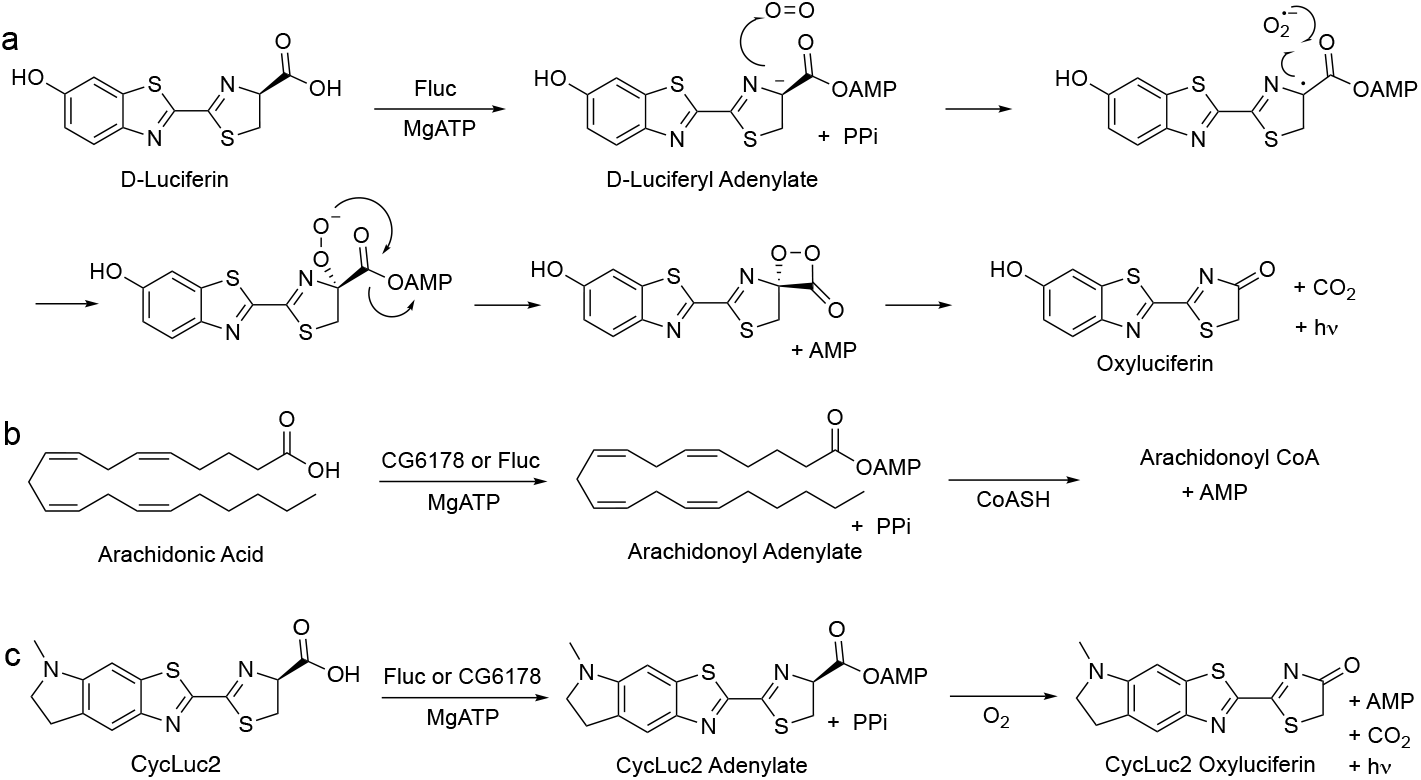
Firefly luciferase and CG6178 both have promiscuous adenylation activity. a) Fluc adenylates D-luciferin, and subsequent reaction with oxygen forms an excited-state oxyluciferin, which emits light. CG6178 is incapable of adenylating D-luciferin and has no luciferase activity. b) CG6178 natively functions as a promiscuous long-chain fatty acyl-CoA synthetase (ACSL), ligating coenzyme A to C8-C20 fatty acids via an adenylated intermediate. Fluc also has ACSL activity. c) Both Fluc and CG6178 have luciferase activity with CycLuc2.

**Figure 2.**
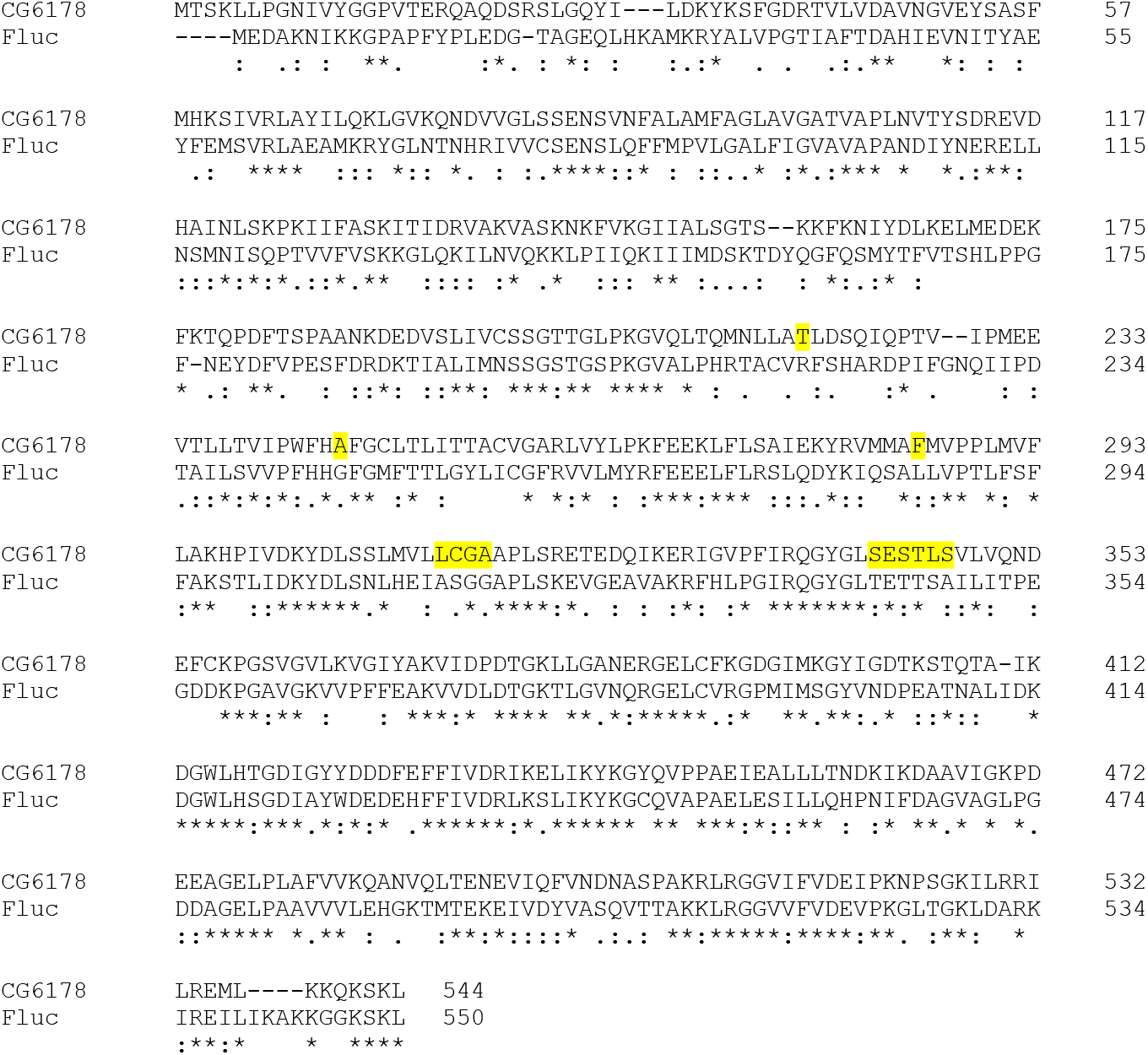
CG6178 shares 41% amino-acid identity with firefly luciferase. *Photinus pyralis* firefly luciferase sequence (UniProt accession number: Q27758) aligned with the CG6178 sequence (UniProt accession number: Q9VCC6) using Clustal under MUSCLE settings.^13^ Amino acid residues within 5Å of the active site selected for mutagenesis are highlighted in yellow.

Here we report the crystal structure of CG6178, which shares the same overall fold as firefly luciferase, but has a steric protrusion occluding its substrate binding site. Through the introduction of mutations to residues that comprise this protrusion, we improved the luciferase activity of this fruit fly enzyme. A mutated variant of CG6178, which we dubbed “FruitFire”, has increased bioluminescent activity with CycLuc2, nascent activity with D-luciferin, and allows bioluminescence imaging in the brains of live mice with CycLuc2-amide.

## Results and Discussion

We first examined whether the latent luciferase activity of CG6178 could be further improved by mutating the active site to more closely resemble that of Fluc. Eleven sets of mutations located within 5Å of the substrate were chosen to replace CG6178 active site residues with the corresponding Fluc residues and broadly evaluate their contribution to luciferase activity. The residues chosen for mutagenesis are highlighted in Figure 2 and the specific sets of mutations used are shown in Supplementary Figure 1. From this limited pool, the mutant enzymes that exhibited the highest luciferase activity were those that swapped residues 342-SESTLS-347 in CG6178 with the corresponding sequence in Fluc (343-TETTSA-348).^14^

We next sought structural guidance for the different enzymatic activity of CG6178 and Fluc and the role of residues 342-347 in CG6178 and 343-348 in FLuc, respectively. CG6178 was crystallized and diffraction data was collected to 2.5Å at the APS 23-ID beamline. Since CG6178 shares 41% identity with Fluc, the structure was determined via molecular replacement using Fluc (PDB ID: 4G36) as a model.^15^ The crystal structure of CG6178 was refined to 2.5 Å resolution with a crystallographic R-value of 0.19 and an R-free value of 0.25 (Supplemental Table 1). 534 out of 544 residues in the CG6178 protein sequence were resolved in the electron density. The first methionine residue, last six residues, and a section from 223-225 were not resolved. The final refined structure shares the same general fold as Fluc (Figure 3), exhibiting an RMSD of 1.45 Å (PDB ID: 4G36).^15^

**Table 1.** *In vitro* Km values for CycLuc2 with CG6178 and mutants. The corresponding amino acid sequence in Fluc is TTSA.

| Enzyme | 344-347 sequence | K<sub>m</sub> (μM) |
| --- | --- | --- |
| CG6178 | STLS | 15 |
| Mutant 1 | STAA | 4.6 |
| Mutant 2 | STAS | 3.8 |
| Mutant 3 (FruitFire) | STSA | 2.1 |
| Mutant 4 | TTSA | 2.4 |

**Figure 3.**
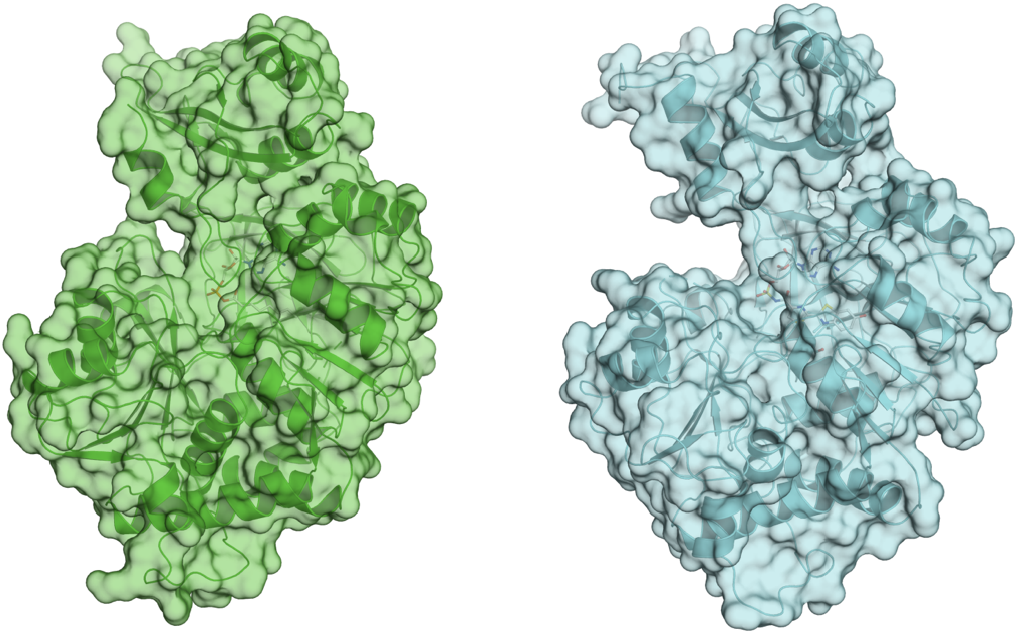
Firefly luciferase and CG6178 share an overall fold. CG6178 (PDB ID: 7KYD, green), and Fluc (PDB ID: 4G36,^15^ cyan). Both enzymes are shown in cartoon and surface representations.

Efforts to co-crystallize CG6178 with either CycLuc2 or D-luciferin were unsuccessful. The addition of ATP dramatically improved crystallization, but surprisingly, density for ATP is not observed within this structure (Figure S2). Rather, the electron density maps are consistent with an adenylated fatty acid (Figure 4, Figure S3). No fatty acids were explicitly added, but CG6178 is known to accept fatty acid substrates ranging in chain length from C8 to C20.^2^ The presence of octanoyl-AMP (C8) and an ethylene glycol in the active site is most consistent with the data (Figure 4, Figure S3).

**Figure 4:**
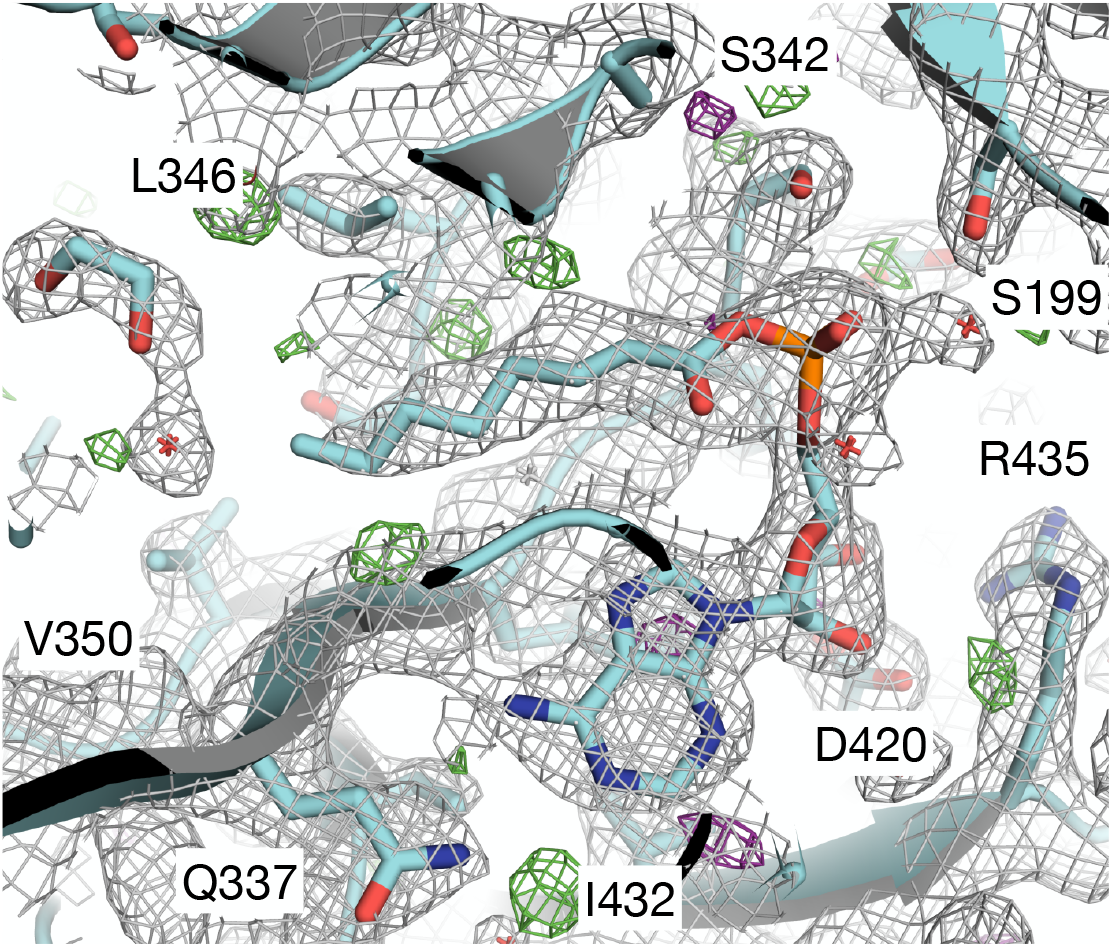
Octanoyl-AMP (C8) represents the best model for the ligand density. CG6178 is shown in cartoon representation. The 2Fo-Fc map (grey, at *σ* = 1) and Fo-Fc difference maps (green *σ* = +3.0 and purple *σ* = −3.0) are shown.

Comparing the active sites of CG6178 and Fluc, residues 342-347 of CG6178 create a protrusion that occludes the binding pocket relative to the corresponding sequence in Fluc (Figure 5), especially due to the Leu346 sidechain. We hypothesized that fatty acid substrates can readily traverse this narrow constriction, whereas the relatively bulky D-luciferin cannot, or at least not in a conformation suitable for attack of its carboxylate on ATP during the adenylation step. Such discrimination is not an issue for CG6178 to perform its native ACSL activity. It is not immediately clear why CycLuc2 would perform better than D-luciferin in this respect; however, CycLuc2 is a more hydrophobic and rigid substrate that may enable some deformation of the active site that is not possible with D-luciferin (Figure 1). Removal of the protrusion in the active site of CG6178 could therefore be beneficial for the luciferase activity of CG6178 with both CycLuc2 and D-luciferin.

**Figure 5:**
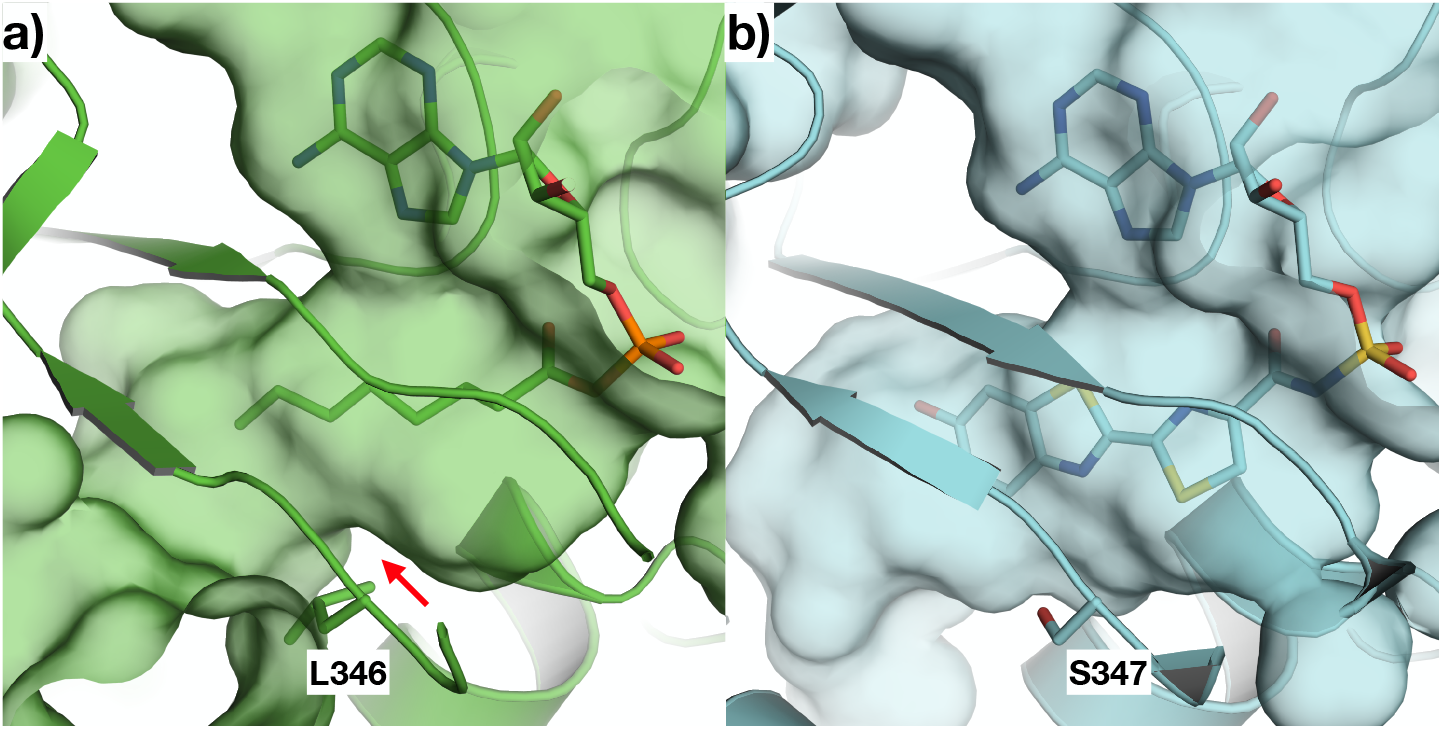
Comparison of the active sites of CG6178 and Fluc. a) CG6178 (PDB: 7KYD; shown in green) and b) Fluc (PDB ID: 4G36^15^; shown in cyan) and were aligned using Schrodinger Maestro^16^, and surfaces shown to compare the substrate binding pocket with (a) octanoyl-AMP and (b) the bisubstrate analog DLSA. The red arrow in (a) shows where CG6178 has a protrusion from Leu 346 that impinges upon the active site.

The sidechain of Leu 346 that forms the bulk of the protrusion in CG6178 aligns with Ser 347 in Fluc (Figure 6). The side-chain hydroxyl group of serine 347 in Fluc makes a hydrogen-bonding interaction with the benzothiazole nitrogen of D-luciferin via an intermediate water molecule.^17^ However, previous work has shown that simply mutating leucine 346 to serine in CG6178 was insufficient to enable CG6178 to efficiently emit light with D-luciferin.^14^ Furthermore, mutation of Ser 347 in firefly luciferase has a detrimental effect on bioluminescence from D-luciferin, but is well tolerated by aminoluciferins like CycLuc2.^9,18^

**Figure 6.**
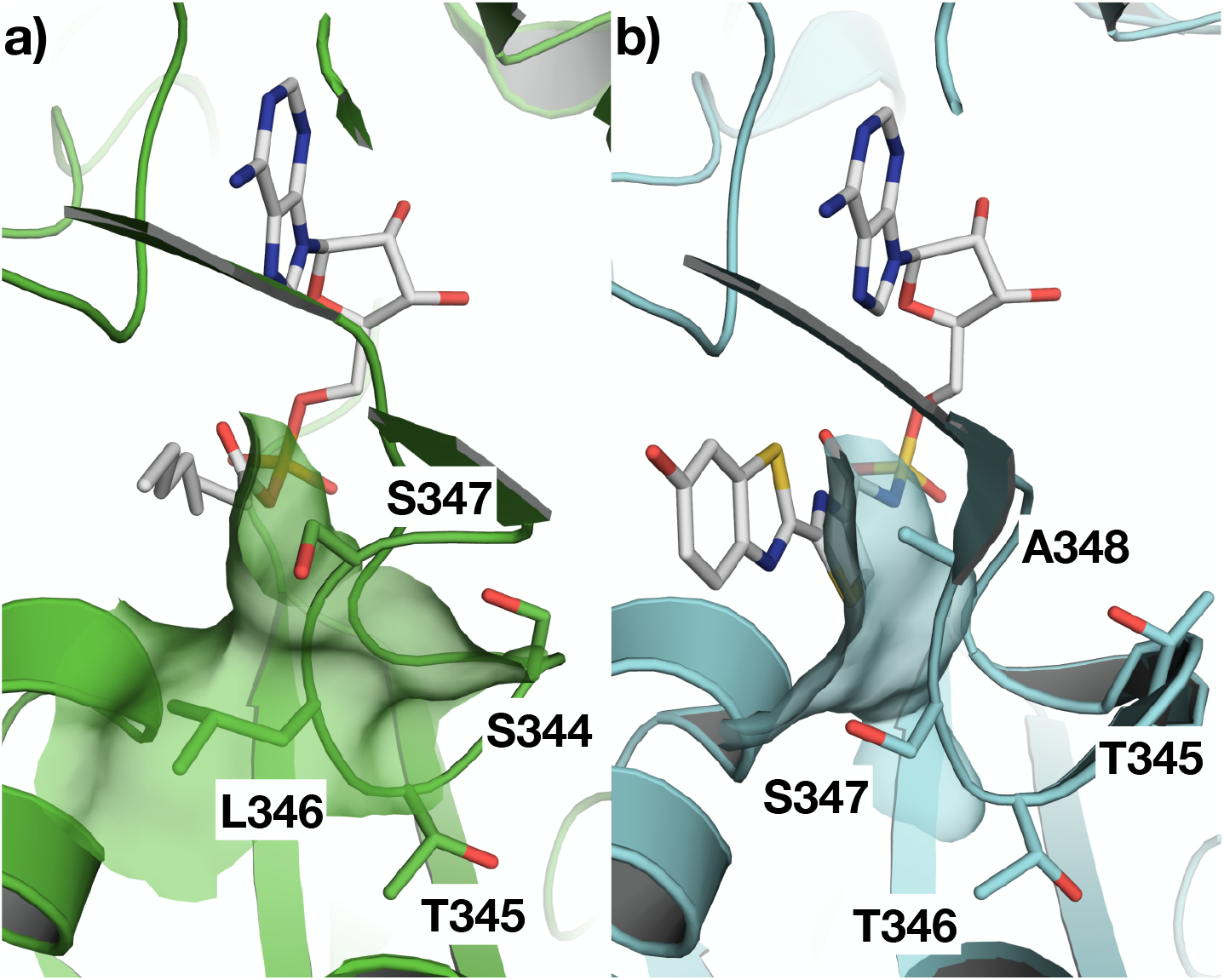
Mutation strategy for improving the luciferase activity of CG6178. The active site of (**a**) CG6178, PDB ID 7KYD; and (**b**) Fluc, PDB ID 4G36^15^ modeled with PyMOL^19^. The residues 344-STLS-347 in CG6178 and 345-TTSA-348 in Fluc are labeled. We mutated CG6178 residues in this loop to better resemble Fluc. Mutant 1: L346A, S347A; Mutant 2: L346A; Mutant 3: L346S, S347A; Mutant 4: S344T, L346S, S347A.

To test our hypothesis that the protrusion created by the 344-STLS-347 loop limits the luciferase activity of CG6178, we generated four mutants of CG6178 that contained mutations to leucine 346 and the flanking residues (Figure 6). These sequences were chosen to reduce the size of the bump in the active site of CG6178 by mutation to alanine, and/or to introduce the residue that occupies this position in firefly luciferase. We found that three of the four CG6178 mutant enzymes improved luciferase activity with CycLuc2 (Figure 7). At 250 μM CycLuc2, emission from mutant 1, 3, and 4 was 2.6, 20.1, and 15.9-fold above that of CG6178, respectively (Figure 7B). Furthermore, these mutants all lowered the Km for CycLuc2 from 15 μM to 2.1-4.6 μM (Table 1), suggesting they would be useful in contexts where substrate access is limiting, such as *in vivo*. Mutant 3, which replaces 344-STLS-347 with 344-STSA-347, exhibited the best performance overall and was renamed FruitFire.

**Figure 7.**
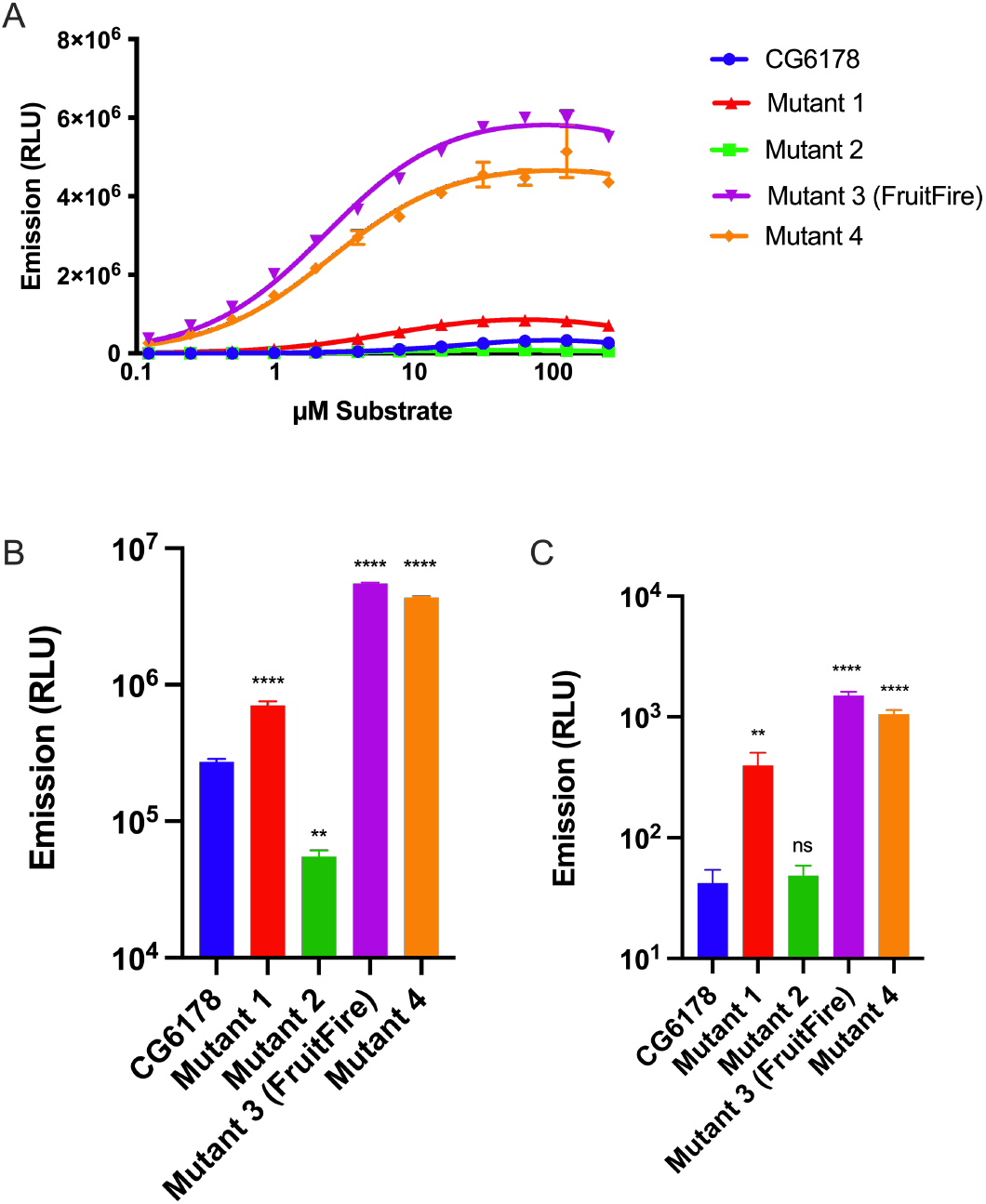
*In vitro* performance of CG6178 mutants. A) Comparison of luciferase activity (40 nM enzyme) with CycLuc2 (0.122 - 250 µM); B) Relative performance with 250 µM CycLuc2; C) Relative performance with 250 µM D-luciferin. Dunnett’s multiple comparison test made to CG6178 (**** = P-value <0.0001, ** = P-value <0.01, ns = not significant).

We next tested whether these mutants would permit any light emission with D-luciferin. Mutant 2, which exhibited poorer luciferase activity than wild-type CG6178 with CycLuc2, did not catalyze light emission from D-luciferin. However, mutants 1, 4, and FruitFire were each able to catalyze weak but measurable light emission with D-luciferin in a dose-dependent manner (Figure S4). At 250 μM D-luciferin, emission from mutant 1, 4, and FruitFire was 9.4, 25.1, and 35.6-fold above background, respectively (Figure 7C), but >1000-fold lower than with CycLuc2 (Figure 7B). Interestingly, mutant 1 enabled light emission from D-luciferin even though it replaces Leu346 with an alanine rather than a serine, and for comparison the simple replacement of leucine with serine only achieved an increase of 3.6-fold over background.^14^ These results suggest that it is not the introduction of a serine at this position per se that critically enables nascent luciferase activity with D-luciferin, but rather the removal of steric protrusion into the active site that includes both leucine 346 and serine 347 of CG6178.

Following the successful improvement of bioluminescent light intensity and lowering of the Km for CycLuc2 through mutagenesis, we next sought to determine whether these results would apply to live mammalian cells. We transfected Chinese hamster ovary (CHO) cells with either Fluc, CG6178, or FruitFire, then incubated them with D-luciferin or CycLuc2 and imaged with a sensitive CCD camera (IVIS-100). Compared to CG6178, FruitFire was brighter and exhibited a lower apparent Km with CycLuc2 (Figure 8).

**Figure 8.**
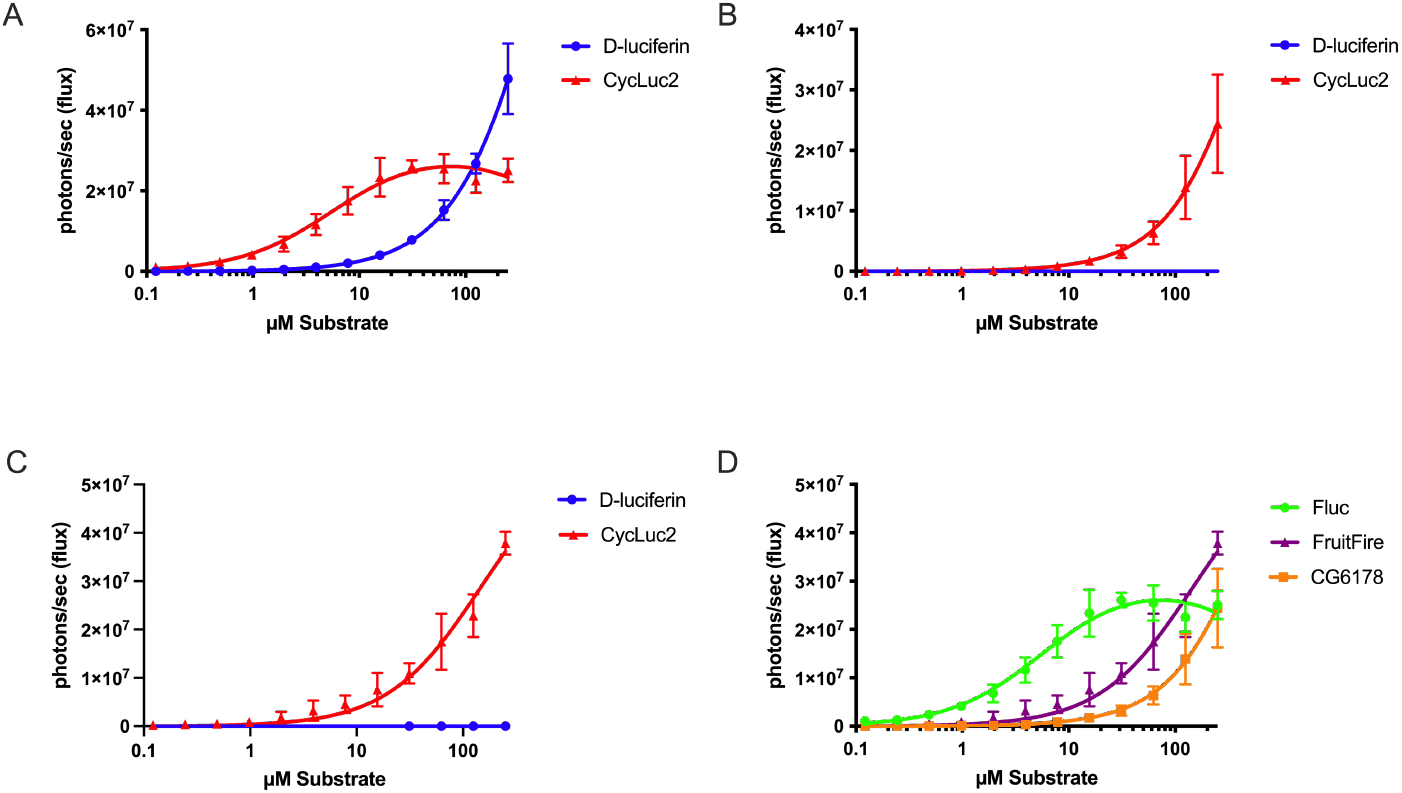
Improved performance of FruitFire over CG6178 in live cells. CHO cells transfected with A) Fluc, B) CG6178, or C) FruitFire were imaged with D-luciferin and CycLuc2; D) relative performance of all three enzymes with CycLuc2.

The lower Km and greater luminescence activity with CycLuc2 compared to CG6178 suggested that FruitFire may be suitable for *in vivo* use. To test this hypothesis, we expressed both enzymes in live mouse brain via stereotactic delivery of adeno-associated viral (AAV) vectors.^18,20^ We have previously used this method to characterize the emission from Fluc and mutants using D-luciferin and CycLuc2.^18,20^ The brain is a challenging tissue to image because it is difficult for the luciferin substrate to cross the blood-brain barrier. Perhaps unsurprisingly, we found that CG6178 expressed in mouse striatum was incapable of light emission above background noise in live animals when treated with either 112 mg/kg D-luciferin or 3.2 mg/kg CycLuc2 (Figure S5). Mice expressing FruitFire similarly lacked bioluminescence when treated with 112 mg/kg D-luciferin or 3.2 mg/kg CycLuc2 (Figure S6). Although both enzymes are capable of luciferase activity in vitro and in live cells using CycLuc2, and FruitFire has a lower Km for CycLuc2 than CG6178, there is insufficient signal coming from the brain in vivo to detect over the background noise of the camera. This may reflect weaker luciferase activity and/or a higher Km for CycLuc2 with CG6178 and FruitFire compared to Fluc (Figure 8D).^9,18^

To help address the higher Km values of CycLuc2 for CG6178 and FruitFire compared to Fluc, and the difficulties in getting luciferin substrates into the brain, we next turned to CycLuc2-amide.^18,21^ CycLuc2-amide is not itself a substrate for Fluc, and must be converted to CycLuc2 by fatty acid amide hydrolase (FAAH), an enzyme highly expressed in the brain, for light emission to occur.^21^ Despite requiring unmasking by this secondary enzymatic activity, CycLuc2-amide greatly improves the access of the substrate to the brain and results in greater light emission from Fluc and Fluc mutants than the equivalent amount of CycLuc2.^18,21,22^

Excitingly, brain bioluminescence was observed when mice expressing FruitFire in the striatum were treated with 0.32 mg/kg CycLuc2-amide (Figure 9). By contrast, mice expressing CG6178 did not yield detectable emission when treated with 0.32 mg/kg CycLuc2-amide (Figure S5C). Together, this suggests that the mutation of two amino acid residues is sufficient to convert the fruit fly enzyme CG6178 into a luciferase capable of bioluminescence imaging in the brain.

**Figure 9.**
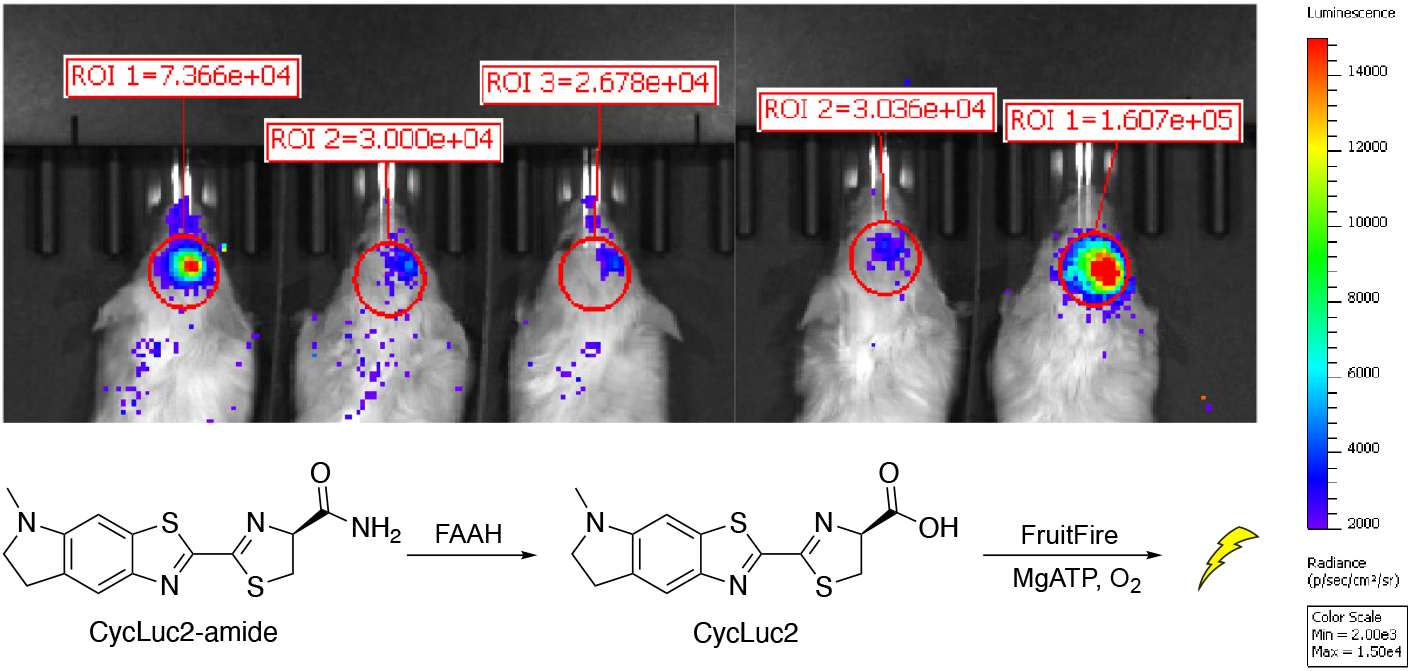
FruitFire emits light in live mouse brain when imaged with CycLuc2-amide. FruitFire was delivered via stereotactic injection of an AAV9 vector into the striatum. Mice were imaged after ip injection with 0.1 mL of 250 µM CycLuc2-amide. Mice 1-3 and mice 4-5 were imaged separately within 30 minutes of each other.

## Conclusions

Many ACSLs from non-luminescent insects share considerable homology to Fluc, and are also generally promiscuous with respect to their fatty acid substrates.^3,4^ Adenylation of a luciferin analogue by an ACSL is proposed to lead to light emission, as such activated intermediates are capable of direct reaction with oxygen and subsequent luminescence.^23^ Currently known examples of latent luciferases among ACSLs from nonluminescent insects are CG6178 and the *Agrypnus binodulus* luciferase-like protein, AbLL.^8,14^ Herein, we sought to elucidate the nature of the substrate discrimination by CG6178 and to improve its luciferase activity. We found that residues 344-347 form a loop that protrudes into the active site. We mutated the leucine and surrounding residues that comprise this bump and found that mutation of just two residues was sufficient to improve light emission by 20-fold, while also lowering the Km for CycLuc2 by ~7-fold. The resulting artificial enzyme FruitFire is capable of bioluminescence imaging in the brain, a tissue that even Fluc with its natural substrate D-luciferin is challenged by.^20,22^ The robust improvement in the luciferase activity of CG6178 via structure-directed mutagenesis highlights the utility of insect ACSLs as platforms for bioluminescent reporter engineering, and more broadly includes enzymes that adenylate carboxylic acid substrates.^4,24,25^ We anticipate that additional mutagenesis can further improve the luciferase activity of insect ACSLs like CG6178, and envision that all ACSLs, regardless of their inherent luciferase activity, can serve as valuable sources of diversity in designing new luciferases.

## Contributions

S.T.A. Jr. performed mouse experiments, purified enzyme and live cell assays, crystallized CG6178, analyzed x-ray data, and prepared the manuscript; J.Z. identified the bound ligand, refined the structure, and made figures; M.F.B. assisted with structural work; C.A.S. advised on structural work; S.C.M. helped devise experiments, analyze data, and prepare the manuscript.

## Acknowledgements

We wish to acknowledge the Kelch lab for collecting the diffraction data, William Royer for guidance in analyzing structural data, and Gordon Lockbaum for assistance in preparing and submitting the structural data to the PDB. This work was funded in part by grants from the National Institutes of Health (EB013270 and DA039961 to S.C.M. and F31 GM116586 to S.T.A. Jr). GM/CA@APS has been funded by the National Cancer Institute (ACB-12002) and the National Institute of General Medical Sciences (AGM-12006, P30GM138396). This research used resources of the Advanced Photon Source, a U.S. Department of Energy (DOE) Office of Science User Facility operated for the DOE Office of Science by Argonne National Laboratory under Contract No. DE-AC02-06CH11357. The Eiger 16M detector at GM/CA-XSD was funded by NIH grant S10 OD012289.

## Materials and Methods

### General

D-luciferin was purchased from GoldBio. CycLuc2 and CycLuc2-amide were synthesized as previously described.^18,26^ Bioluminescence assays were performed on a Xenogen IVIS-100 in the Small Animal Imaging facility. Data acquisition and analysis were performed with Living Image® software. Data are reported as total flux (p/s) for each region of interest (ROI). Data were plotted and analyzed with GraphPad Prism 6.0.

### Plasmid Constructs

Plasmids for bacterial expression were prepared by cloning the target gene into the BamHI-NotI site of a pGEX-6P1 vector. For live cell assays, luc2 and CG6178 were cloned into pcDNA3.1. CG6178 mutants, including FruitFire, were ordered from GenScript.

### Enzyme Expression and Purification

Enzymes were expressed as GST-fusion proteins from pGEX6P-1 vectors. One-liter cultures of BL21-DE3 cells transformed to express the protein of interest were grown at 37 °C until the OD600 was between 0.5-1. The culture was then cooled to room temperature and induced with 0.1 mM IPTG, followed by incubation and shaking at 20 °C overnight. The following day, cells were pelleted at 5000 rpm and flash frozen with liquid nitrogen. The frozen pellets were kept on ice and resuspended in 25 mL lysis buffer (50 mM Tris [pH 7.4], 500mM NaCl, and 0.5% Tween 20) containing 1 mM phenymethylsulfonyl fluoride and disrupted by sonication. Dithiothreitol (DTT) was added at 10 mM, followed by ultracentrifugation at 35k rpm for 1 hour at 4 °C. The supernatant was bound to immobilized glutathione beads (Thermo Scientific) in a Pierce column (Pierce Scientific) for 1 hour at 4 °C. The beads were washed with lysis buffer containing 10 mM DTT, followed by wash buffer (50 mM Tris [pH 8.1], 250 mM NaCl, and 10 mM DTT) and storage buffer (50 mM Tris [pH 7.4], 0.1 mM EDTA, 150 mM NaCl, and 1 mM TCEP). Twenty units of PreScission Protease (GE Healthcare) were added, then the beads were incubated overnight at 4 °C on a Labquake tube rotator (Thermo Scientific) to cleave the GST tag. Protein concentrations were determined on a NanoDrop 100 (ThermoFisher) using the protein mass and molar extinction coefficient method (calculated using the Peptide Property Calculator at: http://biotools.nubic.northwestern.edu/proteincalc.html). For crystallography, CG6178 was further purified using size-exclusion chromatography with a Superdex 75 column (GE Life Sciences). CG6178 was concentrated to 30 mg/mL using cellulose Millipore Amicon® Ultra – 4 Ultracel – 30K centrifugal filters (MilliporeSigma).

### Crystallization of CG6178

To crystallize CG6178, a sitting drop method in Hampton Research Intelli-Plate 48-3 plates was used. After optimized hits following multiple screens, the following method reliably produced crystals within 24 hours: 200 µl of crystallization solution (0.1 M Tris pH 7.5, 0.2 M (NH_4_)_2_SO_4_, 28% PEG 3350, and 10 mM ATP disodium salt) was deposited into the reservoir of each well. Then, 2 µl of the crystallization solution was mixed 1:1 with 2 µl of 30 mg/mL CG6178 in each of 3 sample drop wells within each well. Plates were immediately sealed with Hampton Research ClearSeal Film and stored at 25 °C.

### Crystal Harvesting and Data Collection

Crystals were harvested from their drops with MiTeGen crystal mounts attached to goniometer bases using a MiTeGen wand and deposited immediately into liquid nitrogen. Cryopreserved crystals were transferred to magnetic cryovials and stored in at cryogenic temperatures for transport to the beamline. No cryoprotectant was used. Data was collected at the 23-IDB beamline at the Advanced Photon Source within the Argonne National Laboratory, Chicago, Illinois, U.S.A.

### Data Processing

The crystal diffraction intensities were indexed, integrated, and scaled using HKL3000.^27^ The structure was solved by molecular replacement using PHASER^28^ and using firefly luciferase (PDB 4G36)^15^ as a search model. Model building and refinement were performed using Coot^29^ and PHENIX,^30^ respectively. Ligand models were generated using eLBOW^31^ and the bond length restraints of the ribose ring was adjusted using REEL. The final structure was evaluated with MolProbity^32^ prior to deposition in the PDB (7KYD). To limit the possibility of model bias throughout the refinement process, 5% of the data were reserved for the free R-value calculation.^33^ Structure analysis, superposition and figure generation were done using PyMOL.^19^

### Purified Protein Luminescence Assays – GloMax

50 µL of purified luciferase in enzyme buffer (20 mM Tris [pH 7.4], 0.1 mM EDTA, 1 mM DTT, and 0.8 mg/mL BSA) was added to 50 µL 2X luciferin substrate in substrate buffer (20 mM Tris [pH 7.4], 0.1 mM EDTA, 8 mM MgSO_4_, and 4 mM ATP) in a white 96-well plate (Costar 3912). Luminescence was measured each half second for 1 minute after enzyme addition using a Promega GloMax at a final enzyme concentration of 40 nM and final substrate concentrations ranging from 0.122 µM to 250 µM. Data was exported into Excel for formatting, then pasted into GraphPad Prism for graphing. Data are reported as emission (relative light units, RLU) for each well of the 96-well plate. The values at the final one-minute time point were plotted for each concentration to generate dosage-response curves. Graphs were fit to a sigmoidal dose-response curve by non-linear regression.

### Live Cell Assays

Chinese hamster ovary (CHO) cells and HeLa cells were grown in a CO2 incubator at 37°C with 5% CO2 and were cultured in F-12K Nutrient Mixture (GIBCO) and Dulbecco’s Modified Eagle’s Medium (DMEM) (GIBCO) respectively. Both media were supplemented with 10% fetal bovine serum and 100 U/mL penicillin/streptomycin. CHO cells were transfected with pcDNA3.1 plasmids containing either Fluc, CG6178, or FruitFire. Cells were plated into 96-well black plates (Costar 3915) at 10k cells per well. After 24 hours, cells were transfected with 0.075 μg DNA per well using Lipofectamine 2000 (ThermoFisher). Prior to imaging the transfected cells, a master plate containing the substrates titrated from 250 µM to 0.122 µM over the 12 columns was prepared at sufficient volume to supply substrate to the plates for all three enzymes being assayed. At 24 hours post-transfection, cells were imaged by removing the media containing the Lipofectamine, washing each well with 60 µl Hank’s Balanced Salt Solution (HBSS), aspirating the HBSS, then adding 60 µl per well of the appropriate substrate. After a 3 minute hold, the cells were imaged on a Perkin Elmer IVIS 100. Data collection and Regions of Interest (ROIs) were performed with Living Image Software, then exported to Excel for formatting. The formatted data was plotted using GraphPad Prism and analyzed using nonlinear regression models.

### AAV Vector Construction

Fluc, CG6178, and FruitFire were cloned into the EcoRI-SalI sites of a pAAV-CMV plasmid. AAV-Fluc, AAV-CG6178, and AAV-FruitFire constructs were packaged into AAV9 serotype viral vectors by the Viral Vector Core at the University of Massachusetts Medical School.

### Mice

FVB/N mice were purchased from the Jackson Laboratory. Mouse procedures were approved by the Institutional Animal Care and Use Committee (IACUC) at the University of Massachusetts Medical School (protocol A-2474).

### Striatal Delivery of AAV Vectors

Striatal injections of FVB/NJ mice were performed at 6 weeks of age. Each mouse was anesthetized with isoflurane, then restrained in a stereotactic frame, fitted with an isoflurane nose cone to maintain anesthesia. Mice were kept on a chemical hand warmer wrapped in autoclaved paper towels to maintain body temperature. Hair from the scalp was removed by applying superglue, followed by removing the glued hair with sterilized forceps. The denuded scalps were then washed three times alternately, using 70% ethanol and betadine, using a fresh sterile swab each time. Then, while holding up the scalp with sterilized forceps, a 1 cm anteroposterior incision was made in the middle of the scalp with sterile surgical scissors. The bregma suture was visualized and used as a reference for acquiring the injection site at the lateral edge of the striatum/cortex border (stereotactic coordinates: anterioposterior: +1 mm anterior, mediolateral: +3 mm right lateral, and dorsoventral: −2 mm ventral from bregma). A dental drill wrapped in sterile paper towel containing a sterilized bit was used to drill a hole through the skull at this site, then the needle lowered into the brain at the appropriate depth. After waiting 1 minute, the injection of 1 µl of 2.5e12 GC/ml AAV9 in PBS was delivered at 125 nl/min using a micropump-controlled syringe (NanoFil, World Precision Instruments). After the injection completed, the needle was left in place for one minute before withdrawing. The scalp was sutured using three simple interrupted sutures of dissolvable suture. The mice each then received the following subcutaneously prior to returning them to their cages: 4 mg/kg meloxicam SR, 0.05 mg/kg buprenorphine and 200 µl PBS. The cages remained on heat pads at 37 °C until mice regained their righting reflexes. Bioluminescence imaging was performed at least two weeks after completing the surgeries.

### Preparation of Luciferins for Brain Imaging

D-luciferin (100 mM) and CycLuc2 (2.5 mM) were prepared by dissolving dry compound directly into PBS. CycLuc2-amide was prepared by first dissolving dry compound into DMSO to a concentration of 50 mM, followed by dilution in PBS to a final concentration of 250 µM. Once dissolved, each substrate stock was filtered through a 0.2 µm filter (Millipore Millex GV) prior to injection in mice.

### Brain Imaging

Mice were imaged using the University of Massachusetts Medical School Small Animal Imaging Core using an IVIS 100 imaging system. Mice were induced with isoflurane anesthesia (2% in 1 L/min oxygen), then injected intraperitoneally with 4 µl of luciferin stock in PBS per gram of body mass. Images were acquired as two-minute exposures at 10 minutes after intraperitoneal injection with f/1 and the binning factor set to 8 and analyzed using Living Image software. Regions of interest (ROIs) were drawn around the region covering the brain, and the total flux within each ROI was recorded. ROI sizes were identical across all images.

