## Supplementary Figures 1-6, Table 1 for "FruitFire: a luciferase based on a fruit fly metabolic enzyme"

Table of Contents:

Supplementary Figures 1-6 S2-7

Supplementary Table 1 S8

A

| Mutant # | ASGG (312-315) | F → L (285) | TTSA (344-347) | LCGG (312-315) | TETTSA (342-347) | A → G (245) | T → R (219) |
|---|---|---|---|---|---|---|---|
| 1 | ● | ● | | | | | |
| 2 | ● | | | | | | |
| 3 | ● | | ● | | | | |
| 4 | | | | ● | | | |
| 5 | | | | | ● | ● | ● |
| 6 | | | | | ● | ● | |
| 7 | | | | | ● | | |
| 8 | ● | ● | | | ● | ● | ● |
| 9 | ● | ● | | | ● | ● | |
| 10 | ● | | | | ● | ● | ● |
| 11 | ● | ● | | | ● | | ● |

B

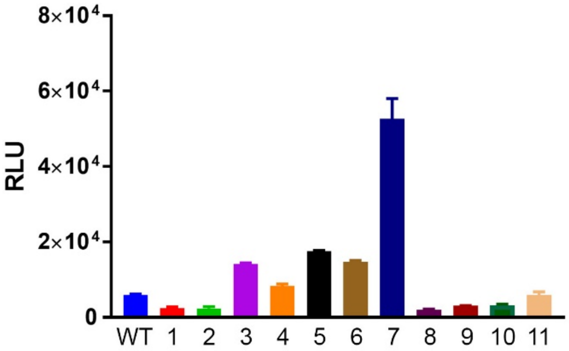


C

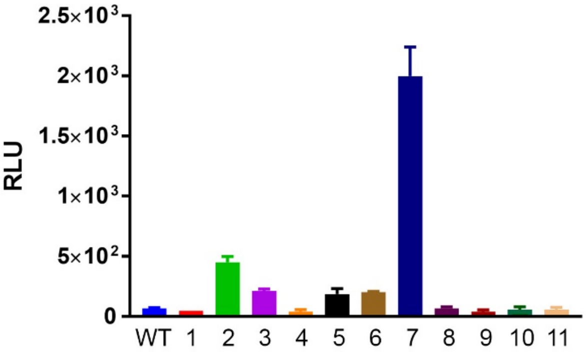


**Supplementary Figure S1. *In vitro* activity of CG6178 active site mutants with CycLuc2 and D-luciferin.** A) Residues in CG6178 highlighted in Figure 2 were replaced with the corresponding Fluc sequence in various combinations as shown. CG6178 and mutants 1-11 (40 nM) were mixed with 250 μM of B) CycLuc2 or C) D-luciferin and light emission was measured with a Promega GloMax.

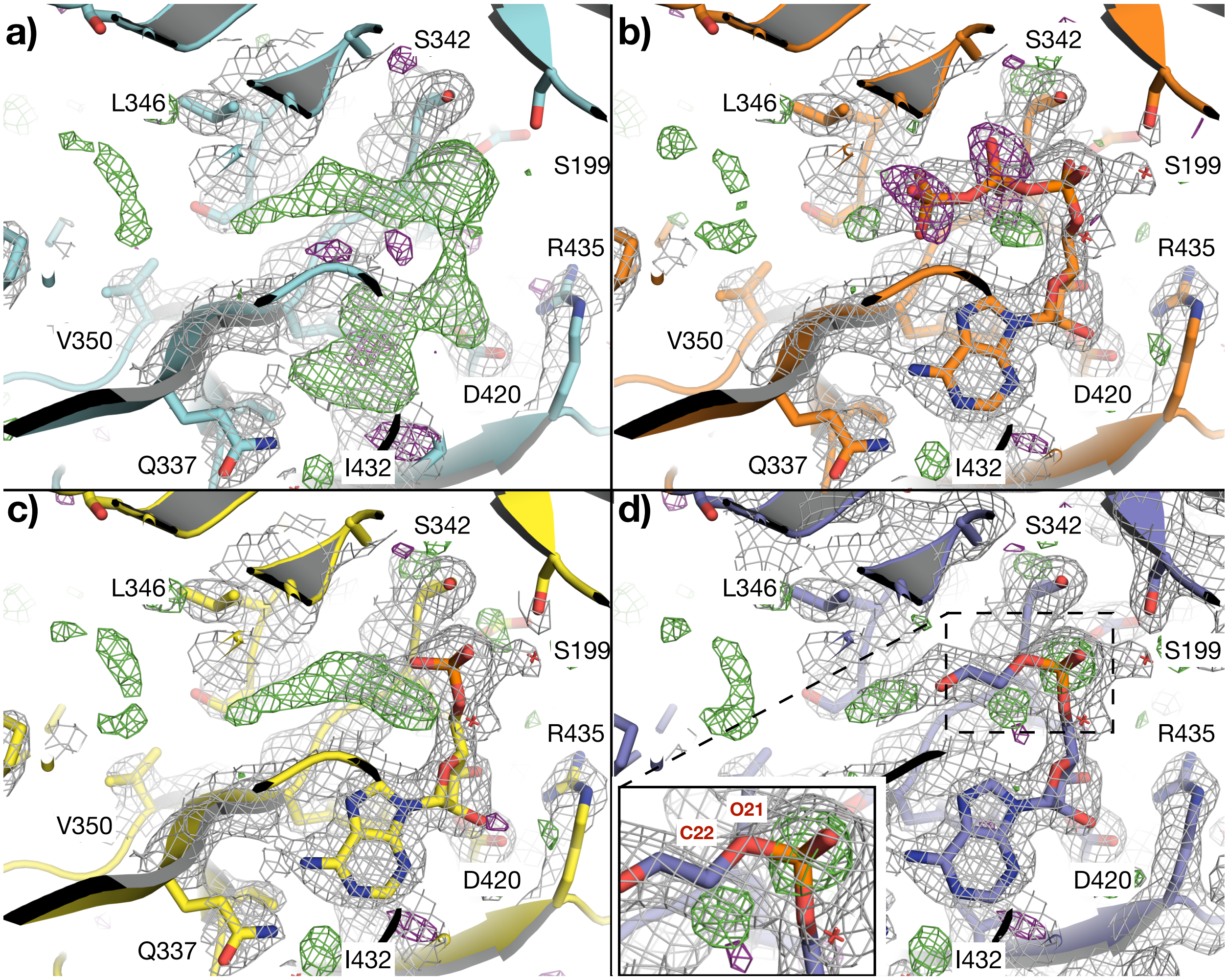


**Supplementary Figure S2**: **ATP, AMP, and an adenylated ethylene glycol are insufficient to satisfy the density in the active site of CG6178**. (**a**) The active site of CG6178 with no ligand modeled. (**b**) ATP, (**c**) AMP, and (**d**) an adenylated ethylene glycol molecule modeled in the ligand electron density. CG6178 is shown in cartoon representation. The 2Fo-Fc map (grey, at $\sigma$ = 1) and Fo-Fc difference maps (green $\sigma$ = +3.0 and purple $\sigma$ = -3.0) are shown.

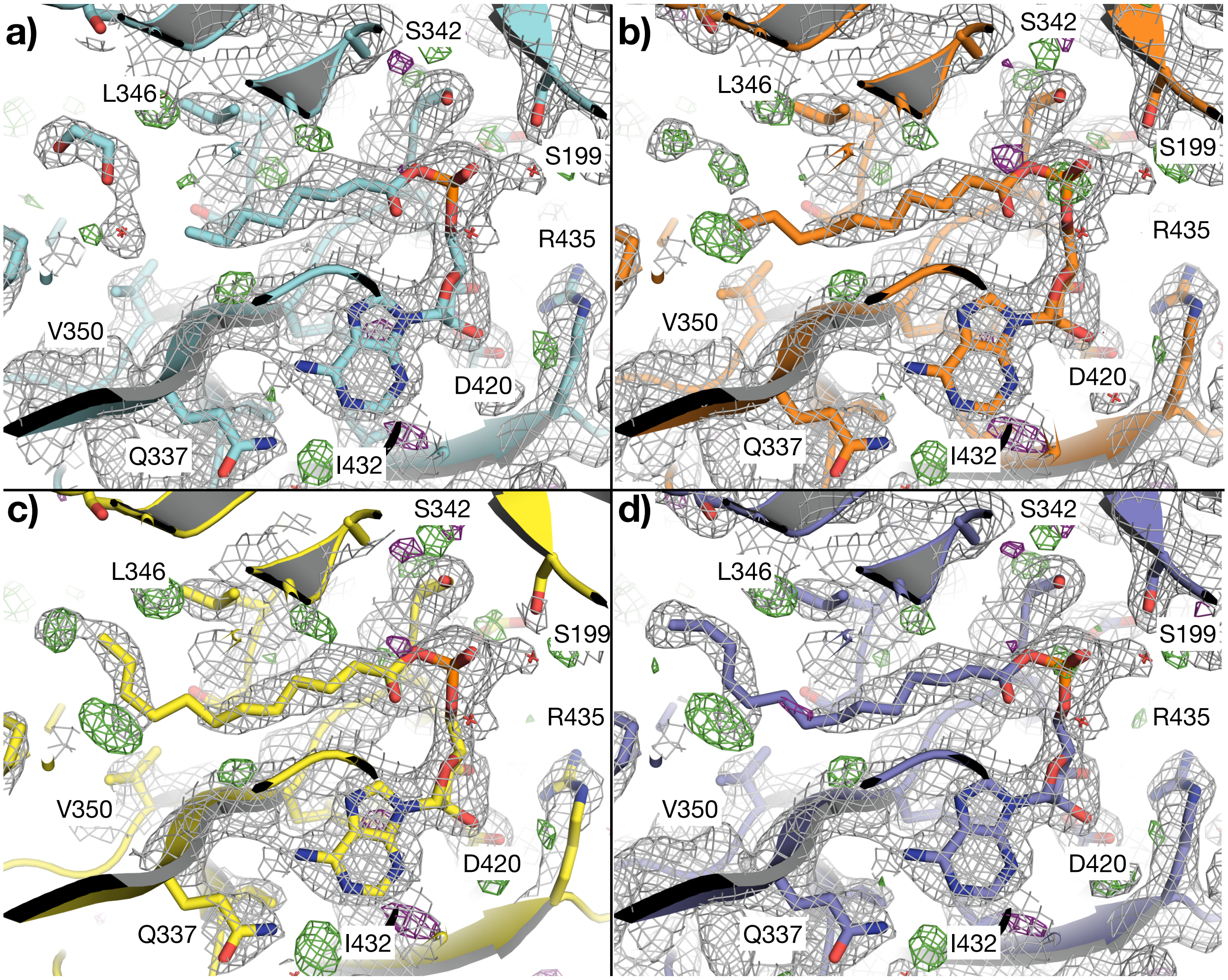


**Supplementary Figure S3**: **Octanoyl-AMP (C8) represents the best model for the ligand density.** (**a**) Octanoyl (C8), (**b**) Decanoyl (C10), (**c**) Dodecanoyl (C12), and (**d**) Myristoyl (C14) AMP esters modeled in the active site of CG6178. CG6178 is shown in cartoon representation. The 2Fo-Fc map (grey, at $\sigma$ = 1) and Fo-Fc difference maps (green $\sigma$ = +3.0 and purple $\sigma$ = -3.0) are shown.

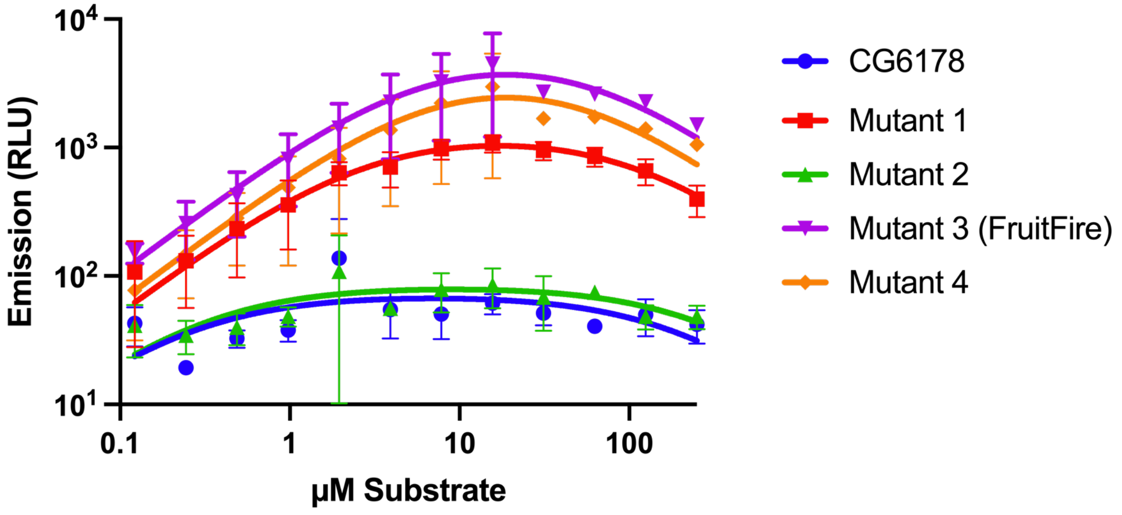


**Supplementary Figure S4.** ***In vitro*** **performance of CG6178 and mutants with D-luciferin (0.122 - 250 µM).** Signal from CG6178 and Mutant 2 is not above background.

A

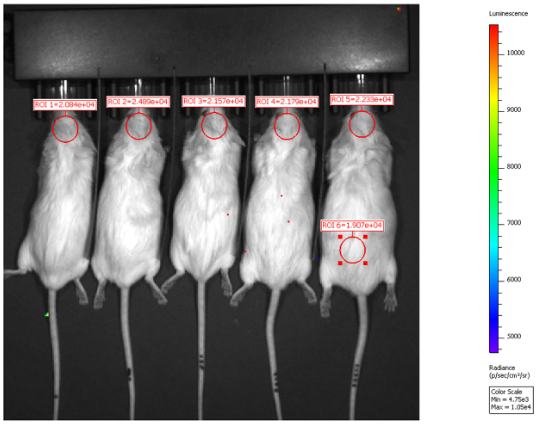


B

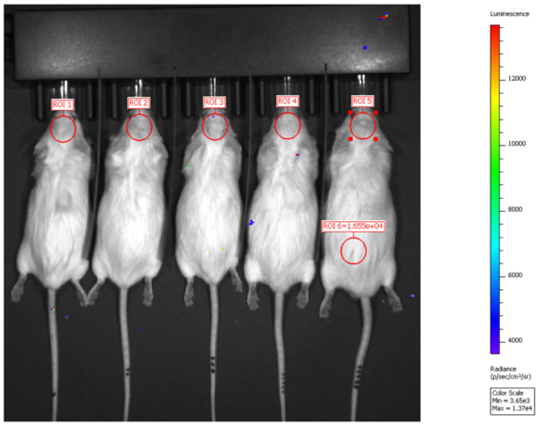


C

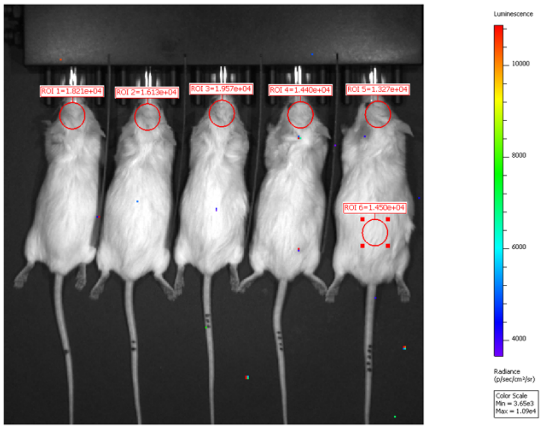


**Supplementary Figure S5. Imaging of mice transduced in the striatum with CG6178.** Mice were imaged after ip injection with A) 0.1 mL of 100 mM D-luciferin, B) 0.1 mL of 2.5 mM CycLuc2, or C) 0.1 mL of 250 µM CycLuc2-amide.

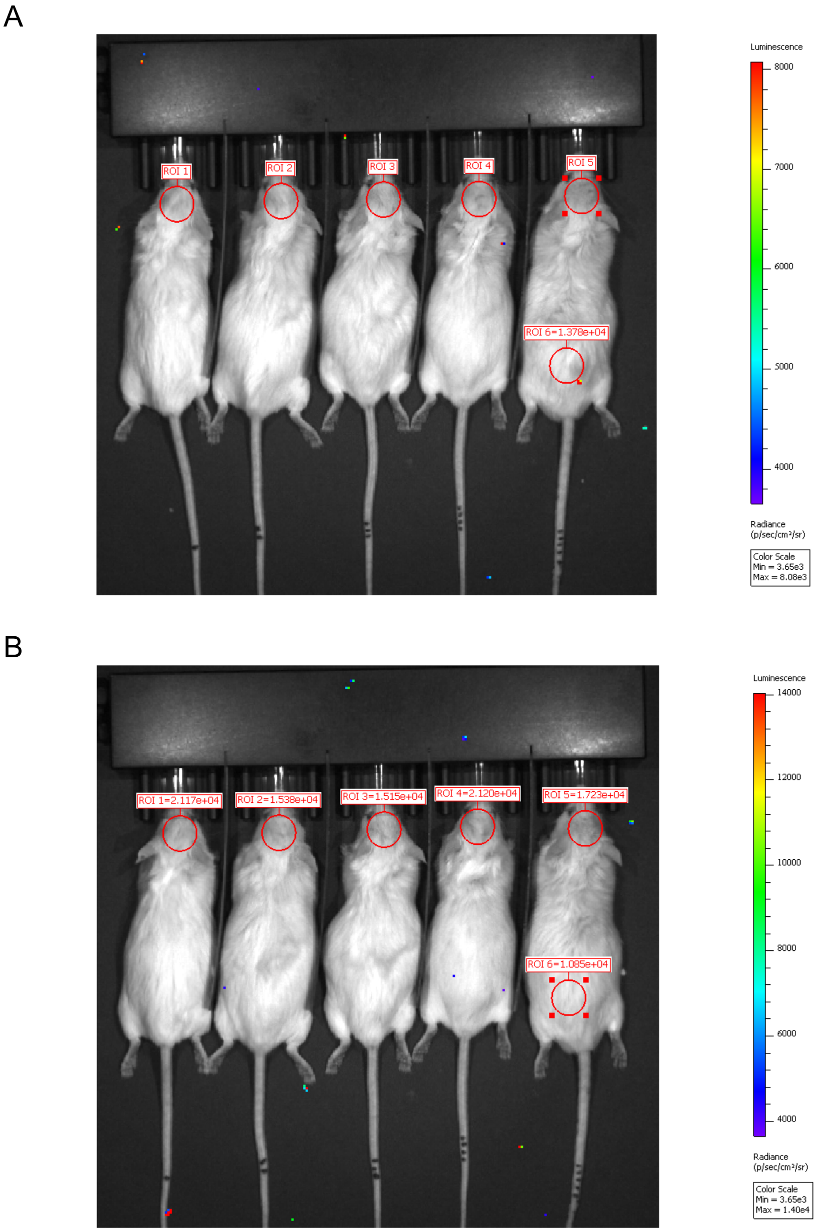


**Supplementary Figure S6. Imaging of mice transduced in the striatum with FruitFire.** Mice were imaged after ip injection with A) 0.1 mL of 100 mM D-luciferin or B) 0.1 mL of 2.5 mM CycLuc2.

**Table S1. Data collection and refinement statistics for CG6178.**

| Data Collection | |
|---|---|
| **Wavelength** | 1.033 |
| **Resolution range** | 30.72 - 2.5 (2.589 - 2.5) |
| **Space group** | P 21 21 21 |
| **Unit cell** | 55.851 77.778 113.315 90 90 90 |
| **Total reflections** | 99870 |
| **Unique reflections** | 17673 (1726) |
| **Multiplicity** | 5.6 |
| **Completeness (%)** | 99.80 (99.37) |
| **Mean I/sigma(I)** | 17.46 (2.95) |
| **Wilson B-factor** | 36.31 |
| **R-merge** | 0.104 (0.671) |
| **Refinement** | |
| **Reflections used in refinement** | 17660 (1724) |
| **Reflections used for R-free** | 885 (87) |
| **R-work** | 0.1949 (0.2515) |
| **R-free** | 0.2527 (0.3272) |
| **Number of non-hydrogen atoms** | 4123 |
| **macromolecules** | 3988 |
| **ligands** | 69 |
| **solvent** | 99 |
| **Protein residues** | 534 |
| **RMS(bonds)** | 0.002 |
| **RMS(angles)** | 0.50 |
| **Ramachandran favored (%)** | 95.53 |
| **Ramachandran allowed (%)** | 4.72 |
| **Ramachandran outliers (%)** | 0.75 |
| **Rotamer outliers (%)** | 1.67 |
| **Clashscore** | 15.16 |
| **Average B-factor** | 38.72 |
| **macromolecules** | 38.72 |
| **ligands** | 42.50 |
| **solvent** | 37.28 |

Statistics for the highest-resolution shell are shown in parentheses.

[b]*R-merge*= $\Sigma \mid I - \langle I \rangle \mid / \Sigma I$, where $I$ = observed intensity, $\langle I \rangle$ = average intensity over symmetry equivalent; values in parentheses are for the highest resolution shell.

[c]RMSD, root mean square deviation.

[d]*R-work* = $\Sigma \| F_o| - |F_c\| / \Sigma|F_o|$.

[e]*R-free* was calculated from 5% of reflections, chosen randomly, which were omitted from the refinement process
